# Loss of a conserved C-terminal region of the *Aspergillus fumigatus* AtrR transcriptional regulator leads to a gene-specific defect in target gene expression

**DOI:** 10.1101/2024.05.22.595332

**Authors:** Shivani Ror, Mark A. Stamnes, W. Scott Moye-Rowley

## Abstract

Treatment of fungal infections associated with the filamentous fungus *Aspergillus fumigatus* is becoming more problematic as this organism is developing resistance to the main chemotherapeutic drug at an increasing rate. Azole drugs represent the current standard-of-care in treatment of aspergillosis with this drug class acting by inhibiting a key step in biosynthesis of the fungal sterol ergosterol. Azole compounds block the activity of the lanosterol α-14 demethylase, encoded by the *cyp51A* gene. A common route of azole resistance involves an increase in transcription of *cyp51A*. This transcriptional increase requires the function of a Zn2Cys6 DNA-binding domain-containing transcription activator protein called AtrR. AtrR was identified through its action as a positive regulator of expression of an ATP-binding cassette transporter (*abcC*/*cdr1B* here called *abcG1*). Using both deletion and alanine scanning mutagenesis, we demonstrate that a conserved C-terminal domain in *A. fumigatus* is required for expression of *abcG1* but dispensable for *cyp51A* transcription. This domain is also found in several other fungal pathogen AtrR homologues consistent with a conserved gene-selective function of this protein segment being conserved. Using RNA-seq, we find that this gene-specific transcriptional defect extends to several other membrane transporter-encoding genes including a second ABC transporter locus. Our data reveal that AtrR uses at least two distinct mechanisms to induce gene expression and that normal susceptibility to azole drugs cannot be provided by maintenance of wild-type expression of the ergosterol biosynthetic pathway when ABC transporter expression is reduced.

**Importance:** *Aspergillus fumigatus* is the primary human filamentous fungal pathogen. The principal chemotherapeutic drug used to control infections associated with *A. fumigatus* are the azole compounds. These drugs are well-tolerated and effective but resistance is emerging at an alarming rate. Most resistance is associated with mutations that lead to overexpression of the azole target enzyme, lanosterol α-14 demethylase, encoded by the *cyp51A* gene. A key regulator of *cyp51A* gene expression is the transcription factor AtrR. Very little is known of the molecular mechanisms underlying the effect of AtrR on gene expression. Here we use deletion and clustered amino acid substitution mutagenesis to map a region of AtrR that confers gene-specific activation on target genes of this transcription factor. This region is highly conserved across AtrR homologues from other pathogenic species arguing that its importance in transcriptional regulation is maintained across evolution.

## Introduction

Fungal infections associated with the pathogen *Aspergillus fumigatus* are a serious threat to health of patients with chronic illnesses especially when the immune system is compromised (1). *A. fumigatus* exists in the environment and easily spreads through its airborne spores. Further complicating the control of aspergillosis is the common appearance of azole resistant *A. fumigatus* isolates that can be found in the environment (2–4). Detailed analyses of these environmental isolates argues that development of azole resistance does not appear to result in any fitness cost, thereby allowing azole resistant isolates to persist and spread via this environmental route (5, 6).

Recognition and analysis of azole resistant *A. fumigatus* isolates led to the identification of two major classes of mutations that trigger this reduction in azole susceptibility. The first and major resistance allele is a compound mutation at the gene encoding the target enzyme of azole drugs, *cyp51A*. This allele consists of two linked changes in *cyp51A*: duplication of a 34 region in the promoter regions (tandem repeat of 34 bp: TR34) along with a replacement of a leucine residue with a histidine residue in the coding sequence of the gene (L98H) (7). A second compound and commonly found allele corresponds to a slightly larger promoter duplication (46 bp: TR46) coupled with two changes in the *cyp51A* coding sequence: Y121F T289A (8, 9). These two different alleles that exhibit reduced azole susceptibility in *A. fumigatus* both lead to strong elevation in the level of *cyp51A* transcription.

Molecular understanding of the mechanism of action of the 34/46 bp repeat region in the *cyp51A* promoter was initially provided by characterization of the SrbA protein, the *A. fumigatus* homologue of mammalian SREBP, a key regulator of cholesterol biosynthetic pathway transcription (10). SrbA was found to bind to an element shared by the 34/46 bp region and is referred to as the sterol response element or SRE (11). Transcriptional activation by SrbA via this SRE leads to induction of *cyp51A* mRNA upon azole treatment (12, 13).

A second factor was found to bind to the 34/46 bp region and during the dentification of a protein designated ABC transporter regulator or AtrR (14). This Zn2Cys6-containing transcription factor is required for normal expression of all forms of the *cyp51A* promoter and is a key positive regulator of multiple *erg* genes, along with SrbA (15). The coordinate control of a key ABC transporter gene (*abcG1*) along with *erg* pathway genes by AtrR provided one of the first examples of the linked regulation of two different means of azole resistance: drug efflux pumps as well as the azole drug target (14).

While the implication of AtrR in control of both *abcG1* and *cyp51A* is clear, there is a deficit in understanding how AtrR acts to stimulate expression of these genes. In this work, we use reverse genetic approaches to identify a region of AtrR that is highly conserved in fungal homologues and critical for gene transcription. Strikingly, this region is essential for normal transcriptional control of *abcG1* but largely dispensable for that of *cyp51A*. These data reveal an unexpected complexity to the coordinate regulation of gene expression by AtrR.

## Results

### Conserved region in AtrR is required for normal azole susceptibility

We aligned the sequences for AtrR homologues from a number of different fungal species and found 3 highly conserved regions. The amino-terminal Zn2Cys6 DNA-binding domain and the associated central regulatory domain are defining characteristics for fungal proteins of this class (16). We also noted a region between residues 855 and 879 of the *A. fumigatus* AtrR that showed strong sequence conservation (Figure 1A). A single amino acid substitution mutation has been identified in this region of the *Sclerotinia homoeocarpa* AtrR homologue that caused the resulting factor to behave as a constitutively active regulator (17). To determine if this conserved region in the *A. fumigatus* AtrR was also involved in gene regulation, we constructed a mutant form of this protein that lacked the entire conserved region. This factor contained an in-frame deletion of residues 855 to 879 and is referred to as Δ855-879 AtrR. All the derivatives of AtrR discussed here are expressed from the wild-type *atrR* promoter and integrated at the normal chromosomal position of this gene. To integrate Δ855-879 *atrR* (and all other derivatives), we placed the *cyc1* termination region downstream of the *atrR* gene as well as a phleomycin resistance gene. We also integrated this selection marker downstream of an otherwise wild-type form of *atrR* to serve as a control for all these experiments.

**Figure 1.**
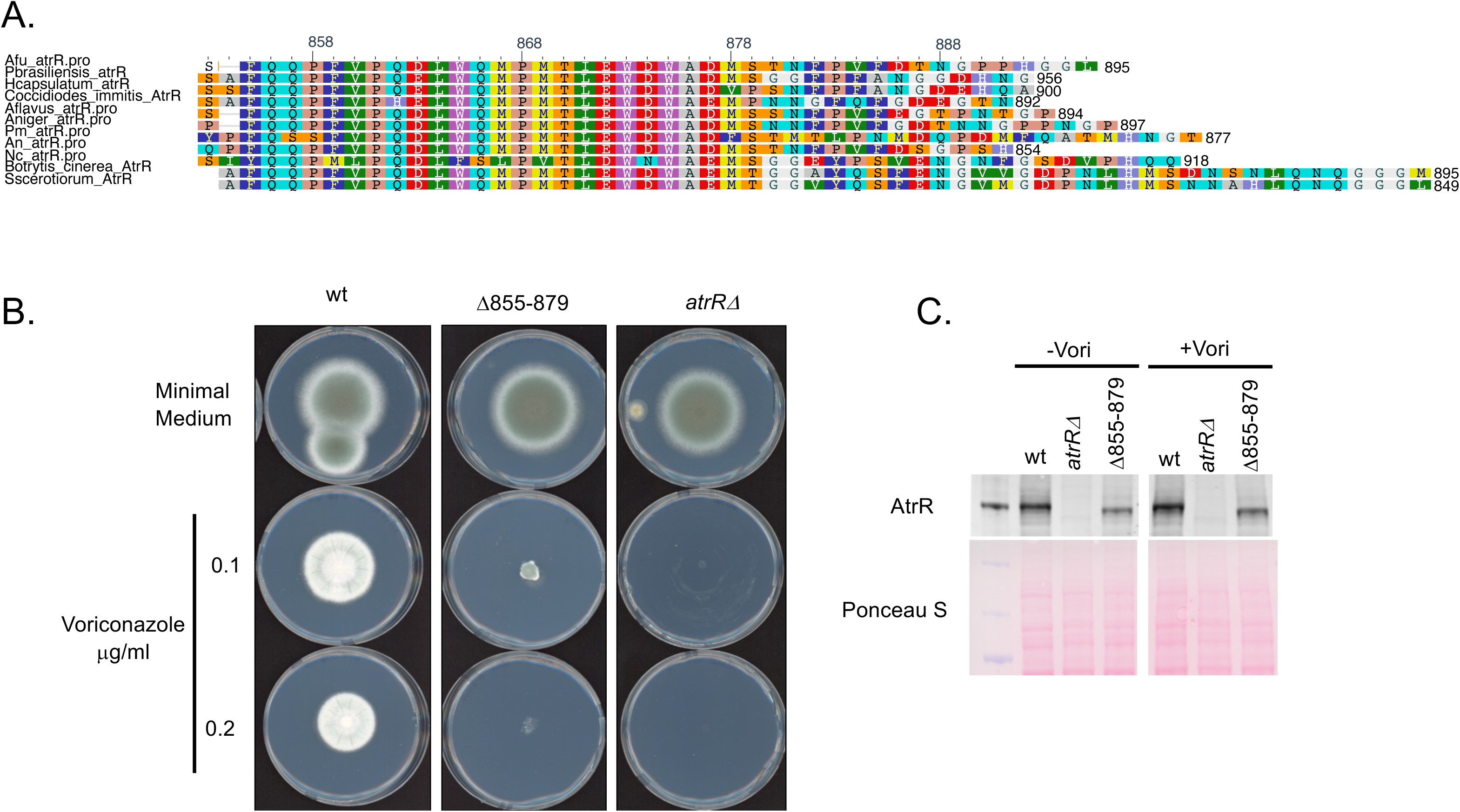
Identification and functional characterization of a conserved C-terminal domain in fungal AtrR proteins. A. Alignments of AtrR proteins from selected fungal species are shown. AtrR homologues from the indicated fungi along with several abbreviated as follows: *A. fumigatus* (Afu), *Paracoccidioides brasiliensis* (Pbrasiliensis), *Histoplasma capsulatum* (Hcapsulatum), *Aspergillus flavus* (Aflavus), *Aspergillus niger* (Aniger), *Penicillium marneffei* (Pm), *Aspergillus nidulans* (An), *Neurospora crassa* (Nc). A plot generated by Multiple Sequence Alignment at NCBI of the conserved region in the AtrR C-termini is shown with the most conserved resides indicated in Rasmol notation. B. Loss of the conserved 855-879 region of *A. fumigatus* AtrR increased voriconazole susceptibility. Isogenic strains containing the indicated forms of AtrR were tested for radial growth in the absence (Minimal Medium) or the presence of different voriconazole concentrations 0.1 or 0.2 μg/ml) C. Loss of the 855-879 region of AtrR does not lead to a large reduction in the steady-state level of the resulting mutant protein. The strains containing the different *atrR* alleles were grown in the presence or absence of voriconazole and whole cell protein extracts prepared. Equal amounts of protein were analyzed by western blotting using an anti-AtrR antiserum (Top panel) and equal loading confirmed by Ponceau S staining of the membrane after protein transfer.

Isogenic wild-type, Δ855-879 *atrR* and the null version of this gene (*atrRΔ*) were placed on minimal medium lacking or containing different concentrations of voriconazole. Plates were incubated at 37°C and radial growth of these strains measured (Figure 1B).

Loss of the 855-879 region of AtrR caused a large increase in voriconazole susceptibility but retained function greater than that of a true null mutant. Based on the observed reduction in radial growth, we estimate that the Δ855-879 AtrR-expressing strain retained 20% of the wild-type activity.

To confirm that this observed increase in voriconazole susceptibility was explained by a loss in activity of the resulting mutant factor, we compared the levels of the wild-type and Δ855-879 AtrR proteins by western blot analyses (Figure 1C). While there is a modest reduction in the steady-state level of the Δ855-879 AtrR factor compared to its wild-type equivalent, significant expression of this mutant protein is retained.

Finally, we examined expression of two of the major AtrR target genes that are both required for normal voriconazole susceptibility: the ABC transporter-encoding *abcG1* locus and the *cyp51A* gene. The strains above were grown in the absence or presence of voriconazole and mRNA levels of these two gene determined by reverse transcription-quantitative PCR (RT-qPCR) using appropriate gene-specific primers (Figure 2).

**Figure 2.**
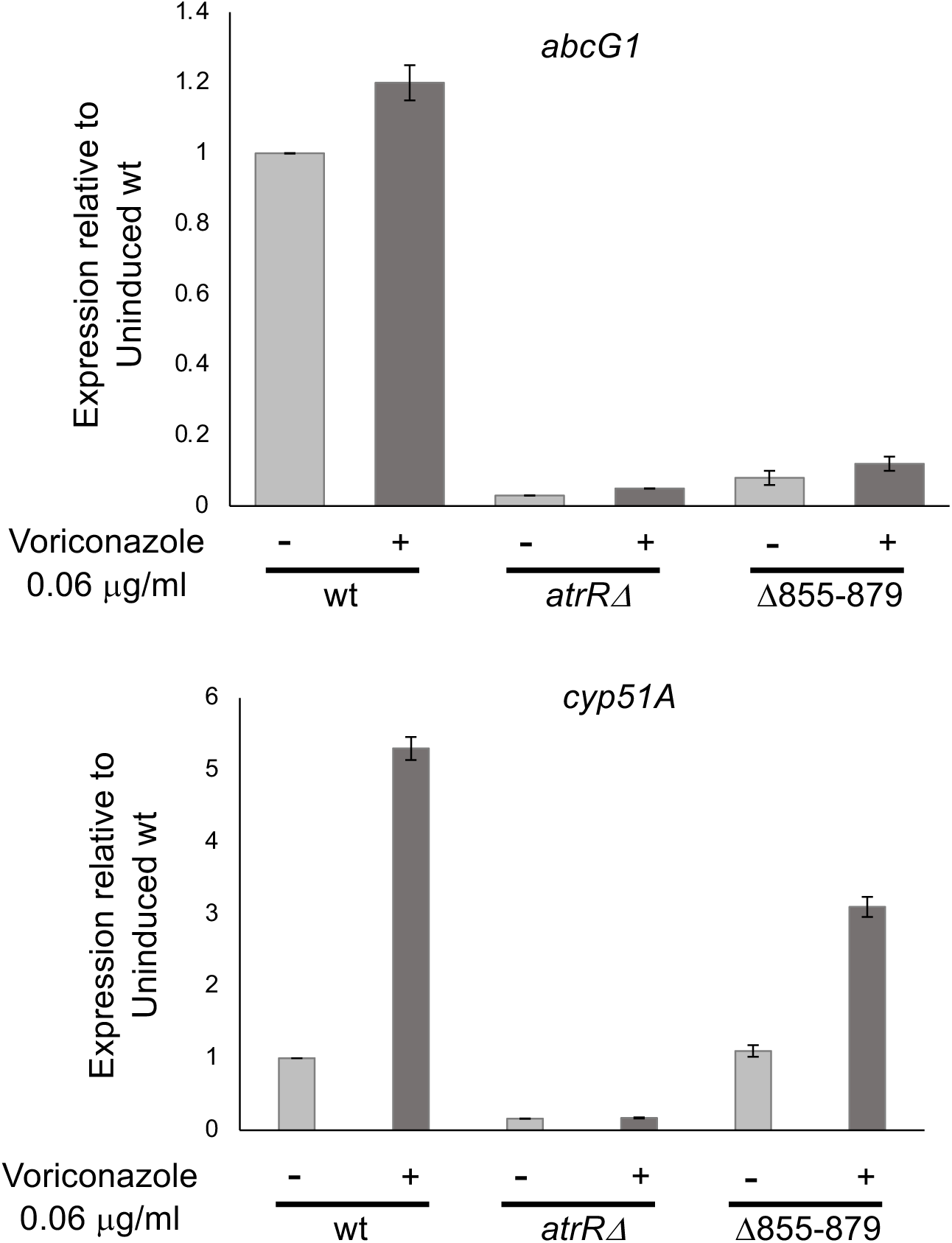
Selective transcriptional defect in the Δ855-879 AtrR protein. Isogenic strains varying at their *atrR* locus were grown 24 hours in the absence (-) or for 16 hours in the absence and then 8 hours in the presence (+) of voriconazole and total RNA prepared. Steady-state levels of mRNA for *abcG1* (top panel) or *cyp51A* (bottom panel) were measured using appropriate primers. Values presented are normalized to the level of expression determined in wild-type cells grown in the absence of voriconazole.

The Δ855-879 AtrR protein exhibited a dramatic reduction in terms of driving expression of *abcG1* as this mutant factor could only produce *abcG1* mRNA levels to <10% those seen in the wild-type strain. Strikingly, Δ855-879 AtrR showed only a minor defect in expression of *cyp51A* that was limited to voriconazole inducible expression.

The presence of the Δ855-879 AtrR induced *cyp51A* expression to 3-fold while the wild-type protein increased transcription to >5-fold. These data provided the first indication that the 855-879 region of AtrR was required for normal transcriptional activation of *abcG1* but only modestly impacted *cyp51A* gene expression. Thus, this mutant AtrR factor exhibited a gene-specific defect in target gene expression.

### Extensive regions of the AtrR C-terminal region are required for normal function

To map the bounds of the transcriptional activation region of AtrR, we prepared a series of C-terminal truncation mutant forms of this factor. The wild-type protein contains 895 amino acids and the 3 truncation mutants correspond to residues 1-854, 1-754 or 1-654. These mutant proteins were all expressed from the normal *atrR* promoter as described above.

Loss of even the final 41 amino acids from the AtrR C-terminus (1–854) led to an increase in voriconazole susceptibility (Figure 3A), best seen at 0.1 μg/ml voriconazole. A similar increased voriconazole susceptibility was seen for the 1-754 form of AtrR while a further truncation leading to expression of 1-654 residues behaved indistinguishably from the *atrRΔ* null mutant strain. Western blot analysis confirmed expression of these mutant proteins although the steady-state level of the 1-654 AtrR derivative was lower than the other factors (Figure 3B). Finally, transcription of *abcG1* and *cyp51A* was assessed under noninduced conditions for all these mutant proteins (Figure 3C). Even truncation mutant with only 41 amino acids removed from the C-terminus of AtrR failed to activate expression of *abcG1* although this mutant factor produced near normal levels of *cyp51A* transcription. The 1-654 form of AtrR was unable to activate expression of either *abcG1* or *cyp51A* and is likely a null mutant form of this factor.

**Figure 3.**
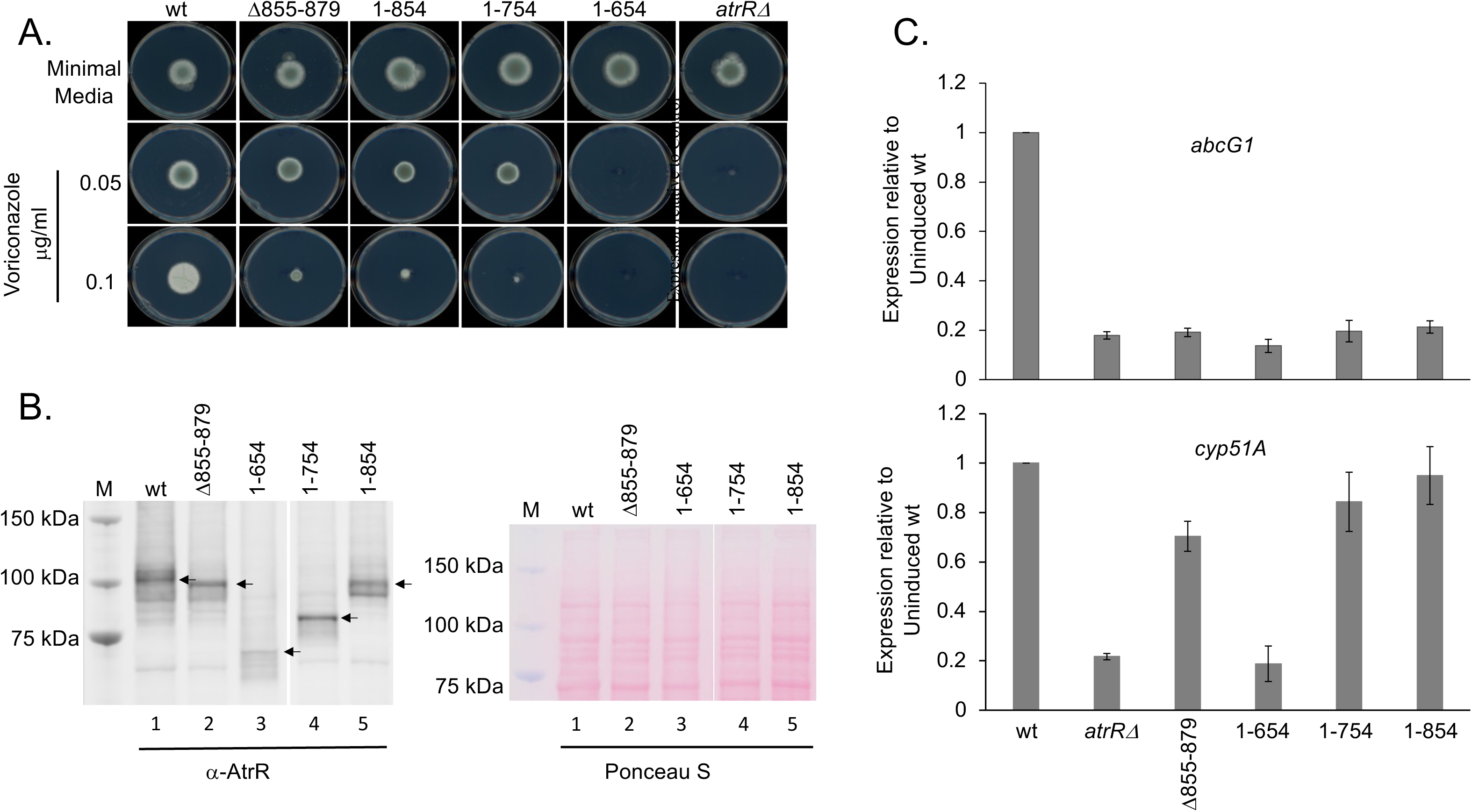
Transactivation domain of AtrR extends to its extreme C-terminus. A. Isogenic strains expressing wild-type (wt) or the indicated forms of AtrR were tested for growth on minimal medium lacking or containing voriconazole at the concentrations listed using the radial growth assay as described above. B. The strains in A were grown for 24 hours in minimal medium and whole cell protein extracts prepared. Equal amounts of protein from each strain were analyzed by western blotting using the anti-AtrR antiserum and equivalent loading confirmed by staining with Ponceau S. The lanes below each panel are identical and correspond to wild-type (Lane 1), Δ855-879 (Lane 2), 1-654 (Lane 3), 1-754 (Lane 4) and 1-854 (Lane 5). M indicates a lane containing molecular mass standards. The arrows denote the major species of AtrR detected. C. Expression levels of *abcG1* and *cyp51A* were determined for each indicated form of AtrR (listed at the bottom for both panels) using RT-qPCR as above.

### Internal deletion derivatives of AtrR map a region required for protein expression

To refine the map of the functional domains of AtrR, we prepared two additional deletion derivatives of the factor. These derivatives were based on the 1-654 AtrR truncation and corresponded to internal deletion in which either residues 755-895 were restored (referred to as Δ655-754 AtrR) or residues 855-895 were restored (Δ655-854). These two factors were analyzed as described for the other truncation mutation proteins above.

Radial growth assays of strains expressing either Δ655-754 or Δ655-854 AtrR on voriconazole-containing media indicated that neither of these derivatives increased the observed function of the 1-654 AtrR protein (Figure 4A). Similarly, expression of both *abcG1* and *cyp51A* remained at the low level driven by the 1-654 AtrR factor (Figure 4B), consistent with these internal deletion mutants behaving as inactive transcriptional regulators.

**Figure 4.**
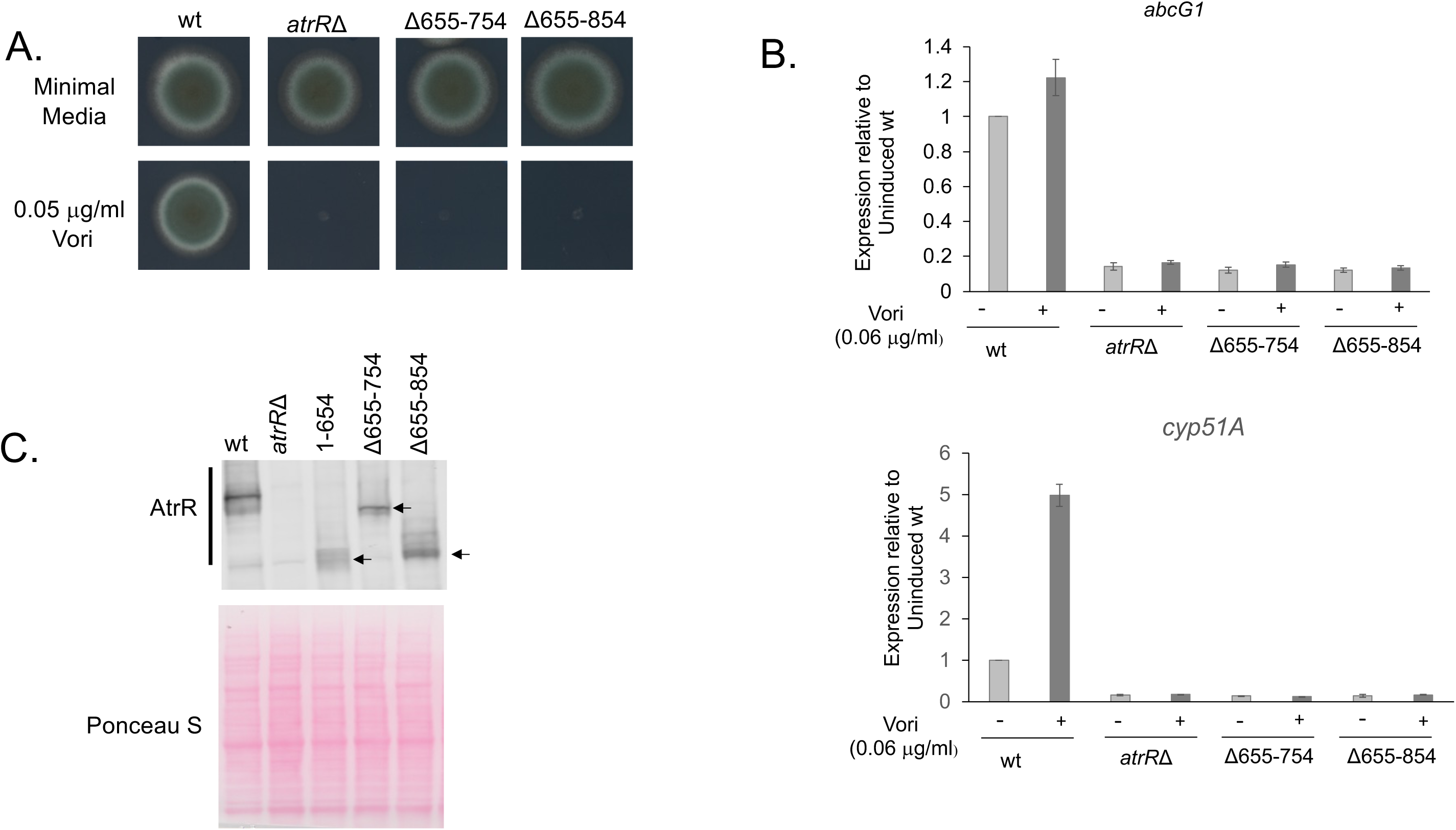
Internal deletions in AtrR identify a region important in expression of the factor. A. The indicated strains were tested in a radial growth assay on either minimal medium alone or containing 0.05 μg/ml voriconazole. B. Expression of *abcG1* and *cyp51A* was assessed by RT-qPCR as above in the presence and absence of voriconazole. C. The strains above along with a control strain expressing the 1-654 C-terminal truncation mutant of AtrR were analyzed by western blotting as above. The estimated location of each AtrR polypeptide chain is indicated by the arrow.

However, comparison of the steady-state levels of both Δ655-754 or Δ655-854 AtrR with the 1-654 AtrR truncation demonstrated that the restoration of as little as 41 amino acids of the C-terminus of AtrR to the 1-654 truncation elevated expression of the resulting mutant protein (Figure 4C). These data support the view that the extreme C-terminal region of AtrR is important to assure normal steady-state levels of this factor.

Measurement of the levels of *atrR* mRNA in these constructs determined that the modest elevation seen in the steady-state mRNAs of these two truncation mutants were not likely to explain the large increase in protein expression (data not shown).

### Alanine-scanning mutagenesis of the 855-879 region of AtrR

To probe the 855-879 segment of AtrR for short stretches of amino acids that could impact function, we prepared triple alanine replacement mutations across this region of AtrR. 6 different mutants were prepared and are designated (for example) FQQ855AAA (where 855 is the first position to be changed to alanine in the AtrR protein chain). The radial growth of these 6 AtrR mutants on voriconazole was tested as above.

The EWD872 and DMS877 AtrR mutants both showed significant reductions in their radial growth on 0.35 μg/ml voriconazole (Figure 5A). The other mutants exhibited either wild-type or marginally elevated radial growth. We also tested these mutants for their effects on expression of both *abcG1* and *cyp51A* (Figure 5B). There was no clear correlation of any of the EWD872 and DMS877 AtrR with reduced gene expression of either gene, although *abcG1* mRNA levels were slightly lower in the case of the DMS877AAA AtrR mutant strain. No alanine scanning mutant exhibited any differential effect on *cyp51A*. We examined levels of both *atrR* mRNA and AtrR protein (Supplemental Figure 1). None of these alanine scanning mutants exhibited a significant alteration from the wild-type in terms of the expression level of the mutant mRNA or protein. Even a range of different 3 amino acid replacements inside the 855-879 region of AtrR cannot mimic the defect seen by loss of this entire region.

**Figure 5.**
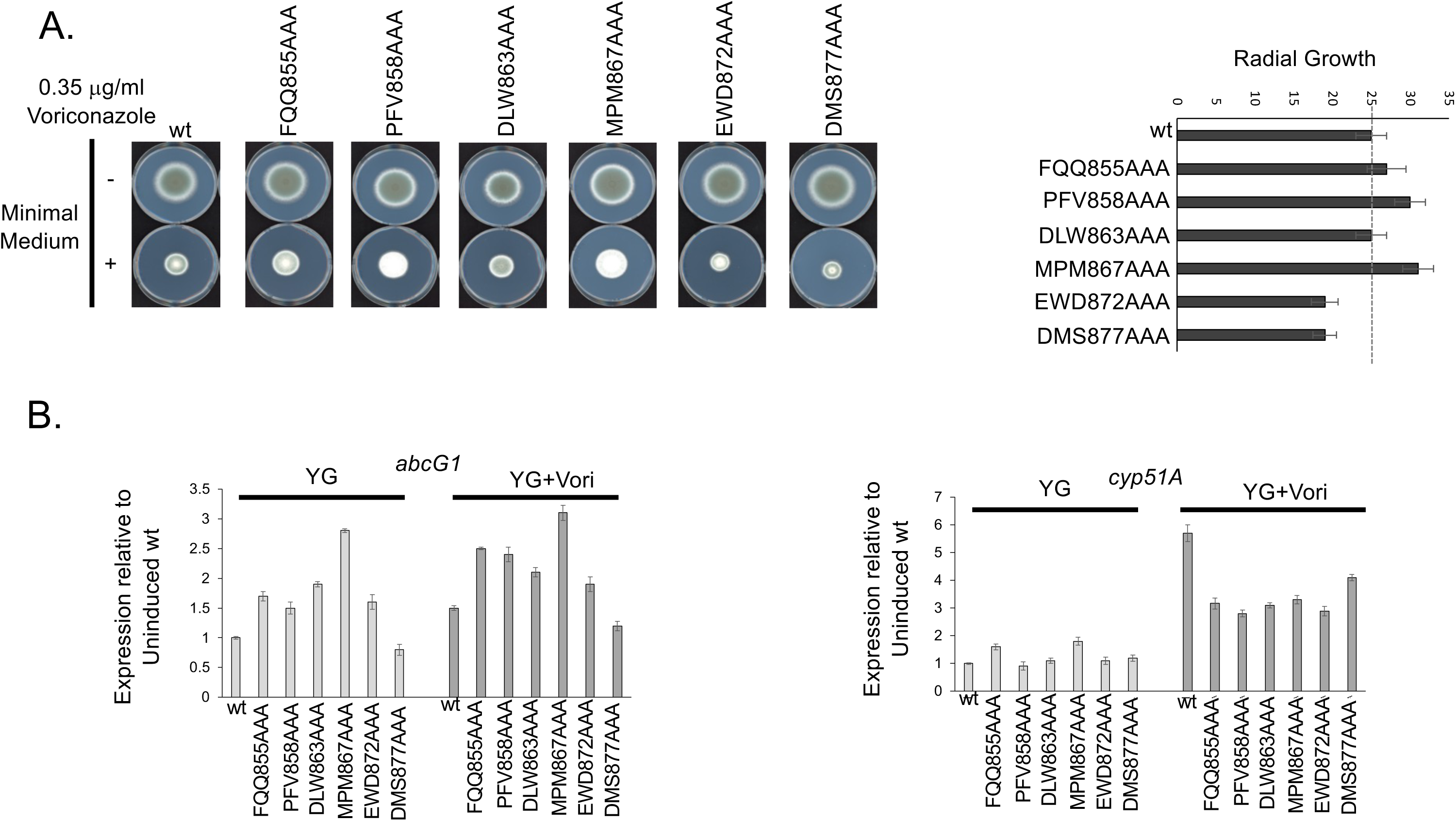
Alanine scanning mutagenesis of the conserved 855-879 region of AtrR. Adjacent three amino acid positions in the conserved 855-879 region of AtrR were replaced with three alanine residues. In each case the 3 amino acids changed are shown along with a number referring to the first position to be changed to alanine. A. The indicated strains were evaluated by the radial growth assay for their ability to grow on minimal medium with or without the indicated concentration of voriconazole. These assays were repeated at least 3 times and diameter of each colony measured. These values are shown in the plot on the right-hand side of the panel. B. Expression of *abcG1* and *cyp51A* was measured by RT-qPCR in the absence and presence of voriconazole treatment.

### Transcriptomic profiling reveals the suite of genes sensitive to loss of the 855-879 region of AtrR

Our initial focus was on the effect of the Δ855-879 AtrR on expression of *abcG1* and *cyp51A*, given the importance of these genes in azole resistance. Our finding with the alanine scanning mutants of AtrR suggests that some degree of voriconazole susceptibility can be caused by defects in AtrR-dependent regulation of loci other than these two genes. To identify all genes that are impacted by loss of the 855-879 region of AtrR, we carried out RNA-seq analyses on isogenic wild-type and Δ855-879 AtrR-expressing strains, both in the presence and absence of voriconazole. Our previous RNA-seq experiments on *atrR* did not test the effect of azole drugs so we also included the *atrRΔ* strain to evaluate the behavior of a true null allele of this transcription factor. Finally, we also used our ChIP-seq data (15) for AtrR to allow us to distinguish between likely direct and indirect target genes.

Since the Δ855-879 AtrR-expressing strain exhibited elevated voriconazole susceptibility, we focused on the overlap between this strain grown on this azole drug compared to the *atrRΔ* null mutant strain, both in the presence and absence of voriconazole. We limited our analysis to genes that significantly (adjusted P-value≤0.05) changed gene expression by a log2≥1 compared to the wild-type strain. These gene sets were overlapped with those known to be positive for AtrR binding in the previous ChIP-seq data (15).

By these criteria, only 37 genes were found to decrease in the Δ855-879 AtrR compared to the wild-type strain in the presence of voriconazole (Figure 6A) with only 9 of these direct targets of AtrR. This finding illustrates the limited transcriptional effect of the Δ855-879 AtrR, although this small number of AtrR-dependent target genes still significantly impacts global azole susceptibility. This small number of responsive genes may be compared to the 1831 genes reduced by the presence of the *atrRΔ* null allele in the presence of voriconazole with 254 of these genes being direct AtrR targets. This is relative to the smaller number of genes in the *atrRΔ* strain grown in the absence of voriconazole in which 524 genes showed reduced expression with 103 direct AtrR target genes. Clearly, the loss of the 855-879 region of AtrR does have an impact on gene regulation but this is very small compared to the sweeping scope of reduction caused by the *atrRΔ* null allele.

**Figure 6.**
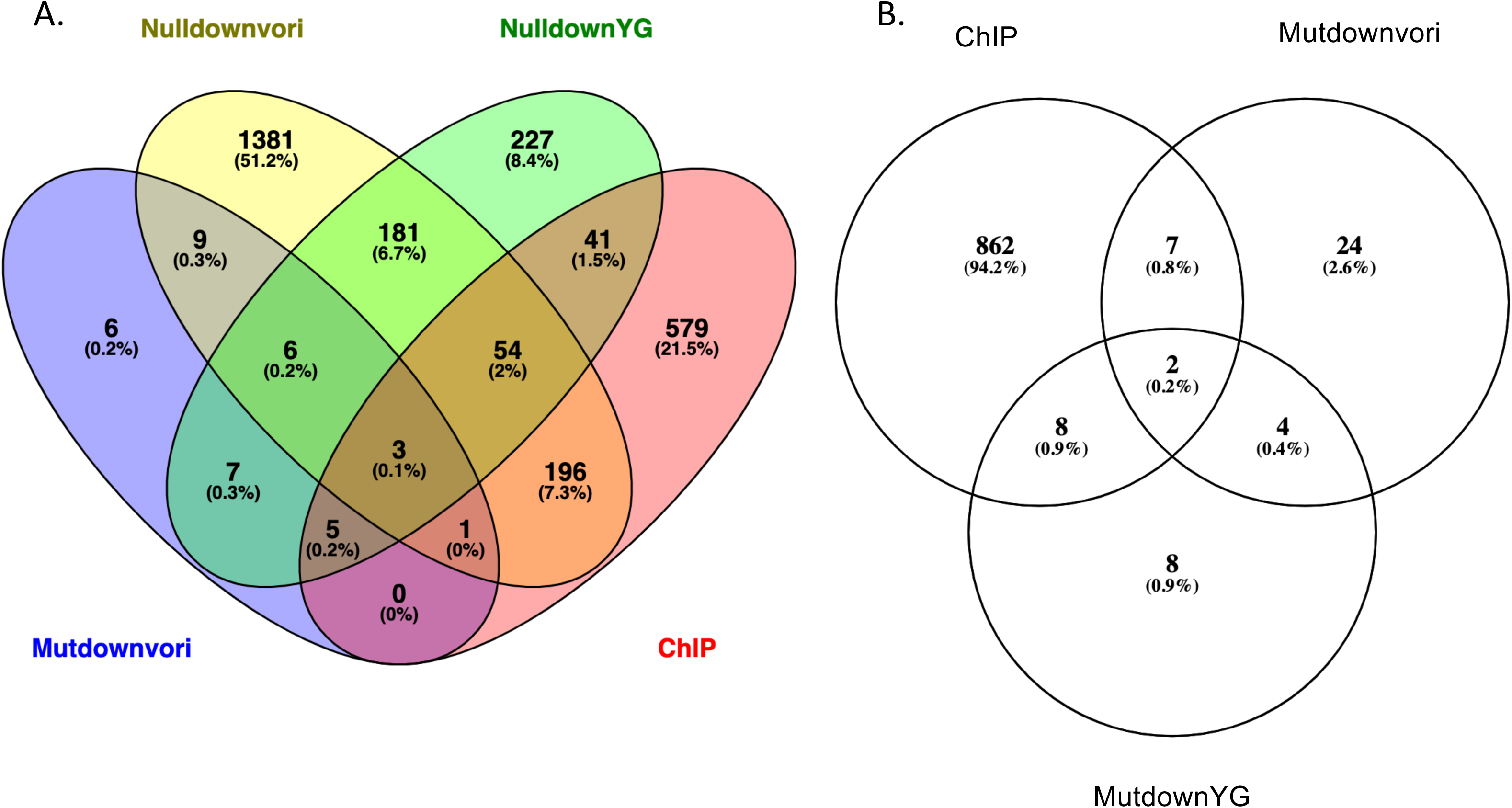
RNA-seq analysis of Δ855-879 AtrR. Venn diagrams of the numbers of overlapping genes from different analyses are shown. All reductions in gene expression are limited to those that are changed significantly (padj<0.05) and a log2≥1. A. Comparison of genes reduced in expression in the *atrRΔ* strain in the absence (NulldownYG) or presence (Nulldownvori) of voriconazole to genes with reduced expression in the Δ855-879 AtrR strain with voriconazole (Mutdownvori). These expression profiles are then assessed for their genes in common with those bound by AtrR via ChIP-seq (ChIP). B. Genes that are reduced in expression in the strain containing the Δ855-879 AtrR protein in the absence (MutdownYG) or presence (Mutdownvori) of voriconazole and that contain AtrR binding sites as measured by ChIP-seq (ChIP).

We next compared the genes seen to be reduced in the Δ855-879 AtrR strain grown in the presence and absence of voriconazole and overlapped these with the direct target set of AtrR-bound genes found by ChIP-seq (Figure 6B). 7 direct target genes were reduced in expression in the presence of voriconazole while 8 were lowered in the Δ855-879 AtrR grown in the absence of voriconazole. 2 genes were lowered under both conditions. Strikingly, 9/17 gene encoded integral membrane proteins that are likely to be transporters (Table 1). Two of these are ABC transporters that have already been implicated in azole resistance: *abcG1* and *mdr1*. The large number of membrane protein-encoding genes responsive to the 855-879 region of AtrR suggests that this domain of the AtrR transcription factor may play an especially critical role in coordinating expression of these membrane components with other AtrR target genes. We also identified two transcription factor-encoding genes that were reduced in the Δ855-879 AtrR strain compared to the wild-type but only during growth on drug-free medium. These two factors, a C6-containing protein (Afu3g05760) and a C2H2-containing protein (ZfpA), may ensure levels of protein prior to drug challenge that are required for normal azole susceptibility. Interestingly, recent experiments have demonstrated that overexpression of ZfpA leads to decreased voriconazole susceptibility, consistent with a role for this protein in this drug phenotype (18).

**Table 1.**
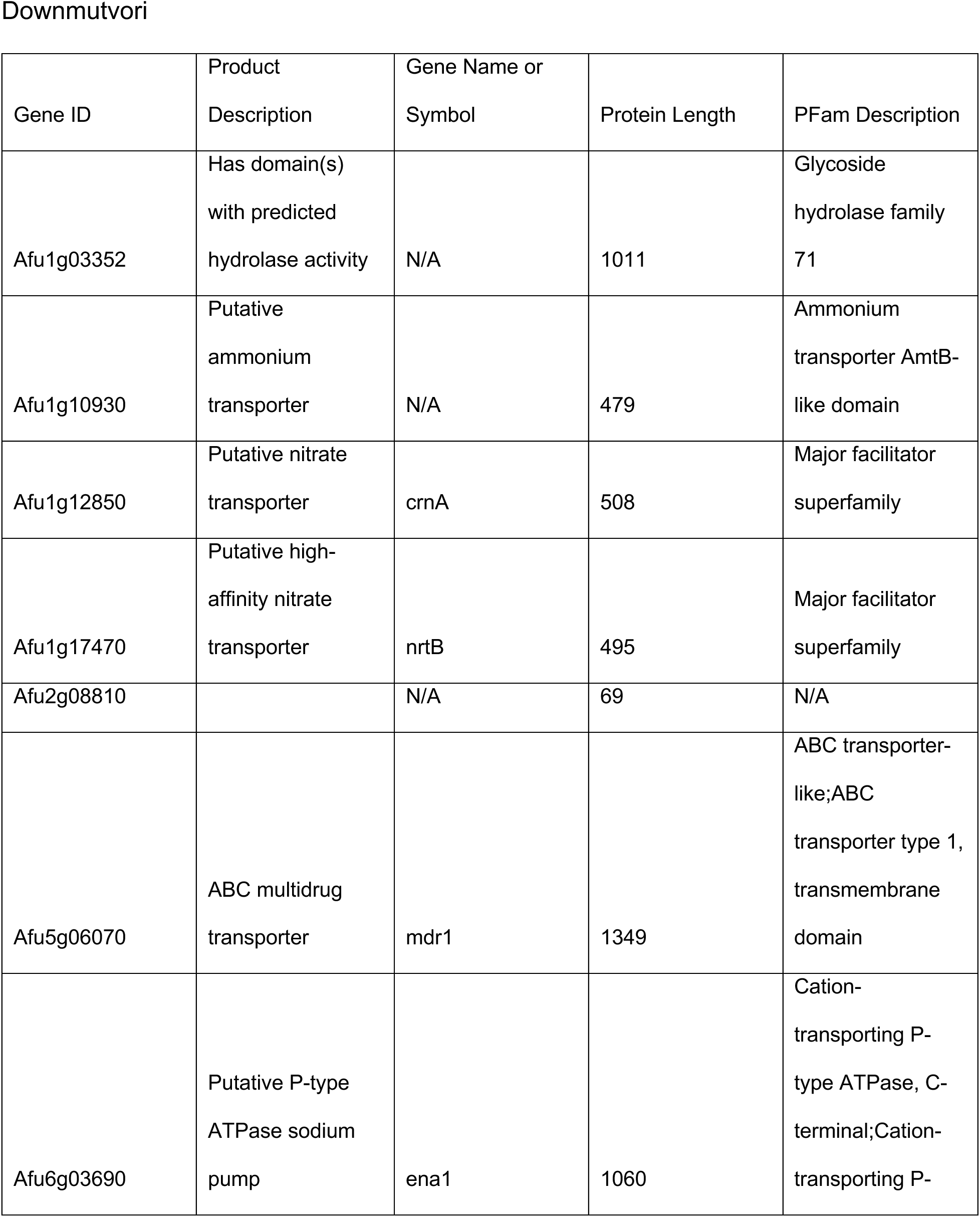
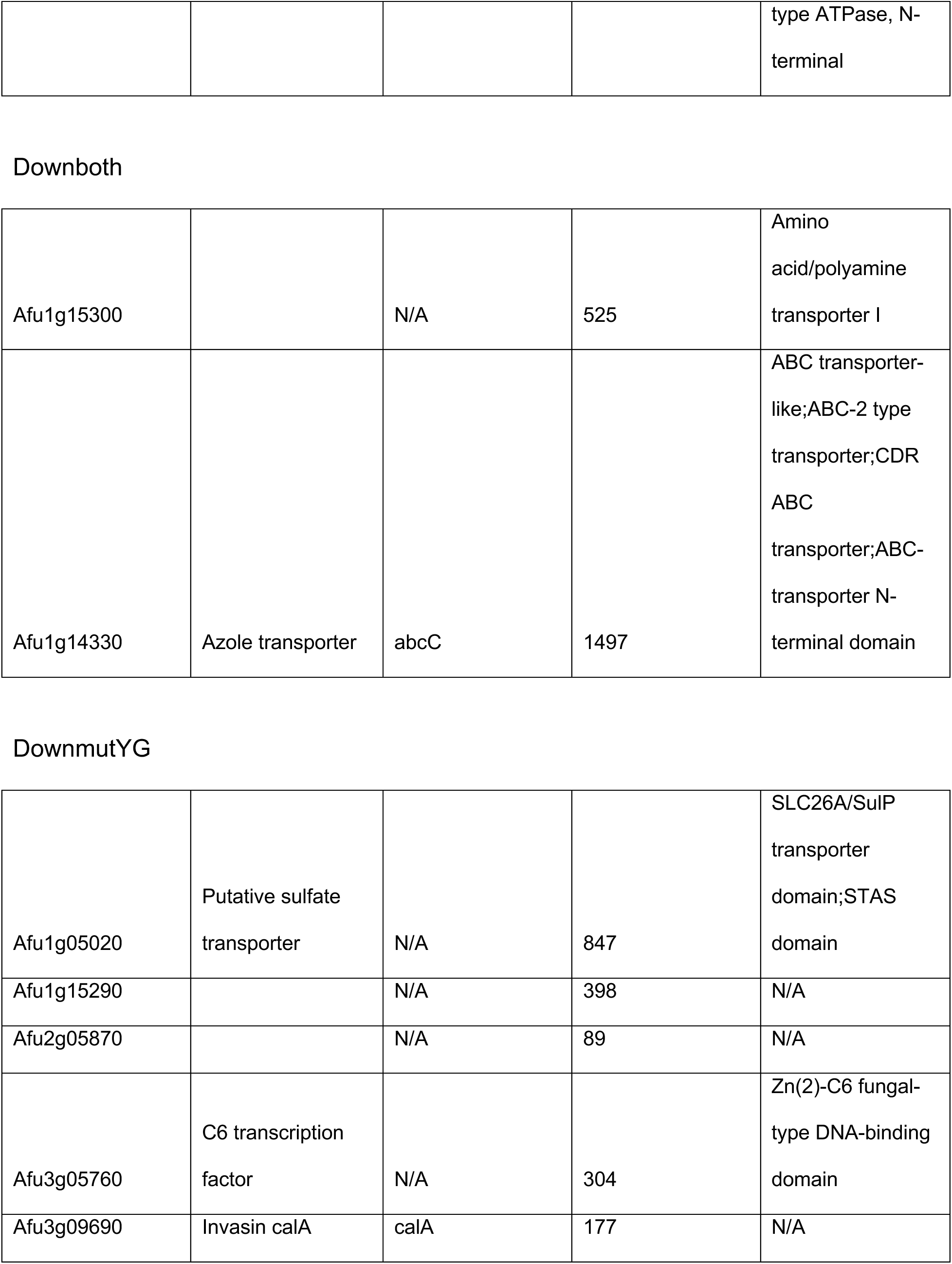
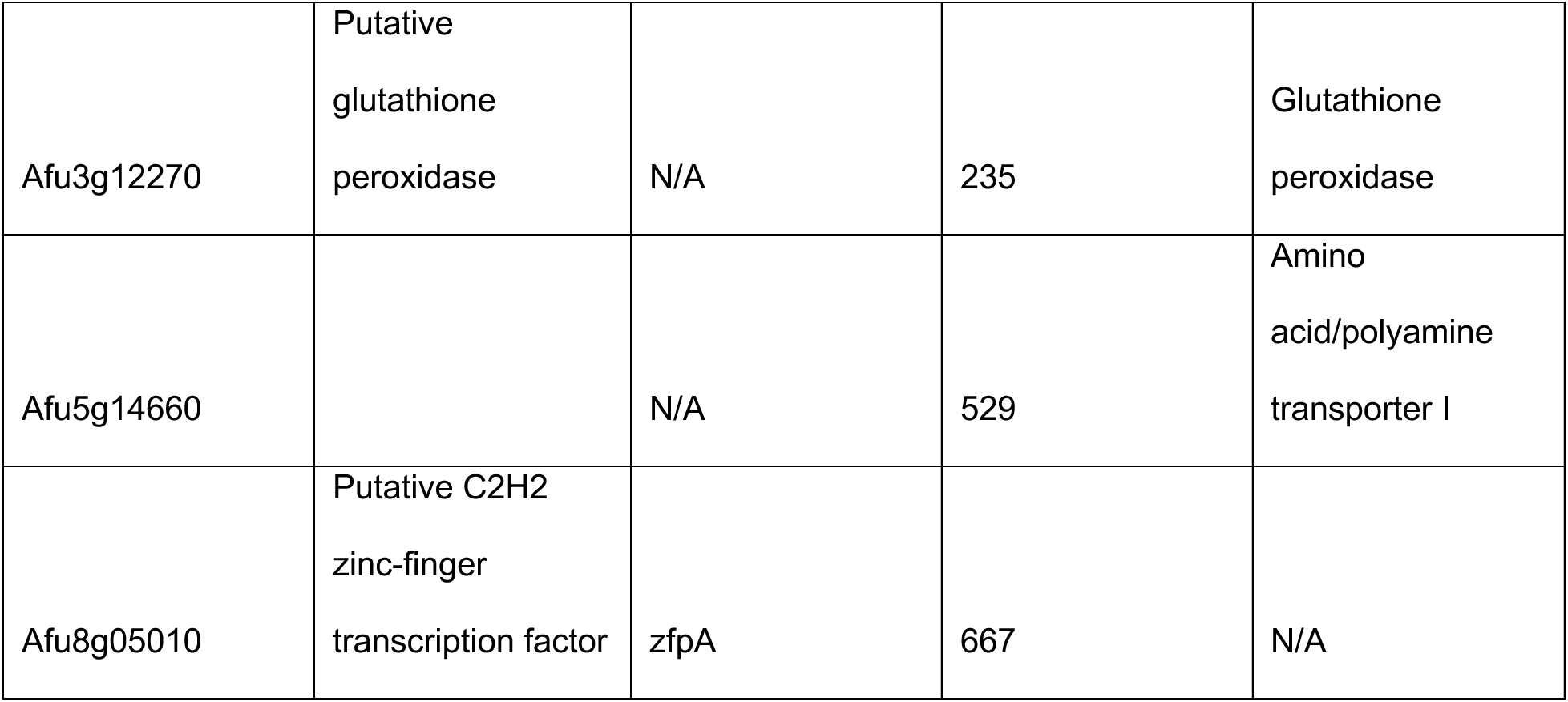
Direct target genes reduced in the presence of Δ855-879 AtrR. . AtrR-bound genes that are significantly lowered by a log2≥1 are listed. Downmutvori, target genes lowered in the presence of voriconazole; Downboth, target genes lowered irrespective of media conditions; DownYG, target genes lowered in the absence of voriconazole.

| Gene ID | Product<br>Description | Gene Name or<br>Symbol | Protein Length | PFam Description |
| --- | --- | --- | --- | --- |
| Afu1g03352 | Has domain(s)<br>with predicted<br>hydrolase activity | N/A | 1011 | Glycoside<br>hydrolase family<br>71 |
| Afu1g10930 | Putative<br>ammonium<br>transporter | N/A | 479 | Ammonium<br>transporter AmtB-like domain |
| Afu1g12850 | Putative nitrate<br>transporter | crnA | 508 | Major facilitator<br>superfamily |
| Afu1g17470 | Putative high-<br>affinity nitrate<br>transporter | nrtB | 495 | Major facilitator<br>superfamily |
| Afu2g08810 |  | N/A | 69 | N/A |
| Afu5g06070 | ABC multidrug<br>transporter | mdr1 | 1349 | ABC transporter-like;ABC<br>transporter type 1,<br>transmembrane<br>domain |
| Afu6g03690 | Putative P-type<br>ATPase sodium<br>pump | ena1 | 1060 | Cation-<br>transporting P-type ATPase, C-terminal;Cation-<br>transporting P- type ATPase, N-terminal |
| Downboth |  |  |  |  |
| Afu1g15300 |  | N/A | 525 | Amino acid/polyamine transporter I |
| Afu1g14330 | Azole transporter | abcC | 1497 | ABC transporter-like;ABC-2 type transporter;CDR ABC transporter;ABC-transporter N-terminal domain |

Table 1. **Direct target genes reduced in the presence of $\Delta 855-879$ AtrR.**
|  |  |  |  |  |
| --- | --- | --- | --- | --- |
| Afu1g05020 | Putative sulfate transporter | N/A | 847 | SLC26A/SulP transporter domain;STAS domain |
| Afu1g15290 |  | N/A | 398 | N/A |
| Afu2g05870 |  | N/A | 89 | N/A |
| Afu3g05760 | C6 transcription factor | N/A | 304 | Zn(2)-C6 fungal-type DNA-binding domain |
| Afu3g09690 | Invasin calA | calA | 177 | N/A |
| Afu3g12270 | Putative<br>glutathione<br>peroxidase | N/A | 235 | Glutathione<br>peroxidase |
| Afu5g14660 |  | N/A | 529 | Amino<br>acid/polyamine<br>transporter I |
| Afu8g05010 | Putative C2H2<br>zinc-finger<br>transcription factor | zfpA | 667 | N/A |

## Discussion

Our understanding of the mechanisms of transcriptional control of gene expression in filamentous fungi is at a relatively early stage compared to other fungal species like the yeast *Saccharomyces cerevisiae* (19). Here we carried out a structure/function analysis on the *A. fumigatus* AtrR transcriptional regulatory protein with the central aim of identifying the transactivation domains of this factor. AtrR is a member of the fungal-specific Zn2Cys6 zinc cluster-containing transcription factors and recognizes DNA via interaction with a linked triplet of CGG residues in its target sites (16). The original fungal AtrR was first isolated from *Aspergillus oryzae* on the basis of its effect on reducing azole susceptibility when overexpressed (14). Later work showed the *A. fumigatus* AtrR acted as a positive regulator of the expression of both *cyp51A* and *abcG1*, two key determinants in the control of azole susceptibility (14, 15).

This work demonstrates that the carboxy-terminus of AtrR is crucial for activation of expression of both *abcG1* and *cyp51A*. We initiated our characterization of the C-terminus of AtrR by deletion mutagenesis of a region that is conserved across fungal AtrR proteins. Additionally, a point mutant form of the AtrR protein from *Sclerotinia homeocarpa* was found that exhibited a large decrease in azole susceptibility coupled with increased expression of the *S. homeocarpa cyp51A* homologue and a number of membrane transporter proteins (17). This mutation is thought to convert the *S. homeocarpa* AtrR into a hyperactive transcription factor. This behavior is similar to that seen for the *Candida glabrata* Pdr1 protein in which many different mutations have been mapped to the C-terminus of this protein that cause the resulting mutant Pdr1 protein to behave as a constitutively active regulator of gene expression (20). We are currently analyzing the behavior of an analogous mutant form of *A. fumigatus* AtrR to determine if similar hyperactive characteristics are seen.

The effect of the deletion of the conserved 855-879 region in AtrR was quite striking as this mutant protein still showed normal gene regulation for the overwhelming majority of AtrR target genes. We only found 17 genes that were both bound by AtrR and exhibited reduced gene expression (Table 1). When a similar analysis is done for the transcriptome of the *atrRΔ* strain challenged with voriconazole, 254 AtrR direct target genes are reduced by loss of this transcription factor. Comparison of the effect of voriconazole on the genome of strains expressing the Δ855-879 AtrR protein relative to the wild-type factor showed that fewer than 500 total genes were significantly altered (either increased or decreased) by log2≥1, indicating that the vast majority of voriconazole-responsive genes (nearly 4000 in the *atrRΔ* null) behave normally in the presence of this mutant AtrR derivative. This limited effect of the Δ855-879 AtrR illustrates the gene-selective defect caused by loss of this region of AtrR.

While the deletion of the 855-879 domain of AtrR clearly showed transcriptional defects, albeit to a limited number of genes, our attempts to identify shorter sequence elements inside this region by alanine scanning mutagenesis yielded less insight. Most of these triple alanine alterations failed to dramatically alter the voriconazole susceptibility of the resulting mutant strain or expression of the two key azole target genes we examined (*abcG1* and *cyp51A*). We suggest this behavior has two likely explanations. First, this could provide evidence for redundancy inside the 855-879 region of AtrR since replacement of a number of different amino acids with alanine residues is well-tolerated, we may only be impacting a single interaction with these substitutions while deletion of the entire domain eliminates all interactions. We will test this by combining multiple alanine substitutions in hopes of mimicking the effect of the 855-879 deletion. A second possibility is that the region responsible for the effect we observed in the presence of the Δ855-879 AtrR overlaps with the deleted region. The fact that the triple alanine scanning mutations made at the end of the 855-879 region (EWD872 and DMS877AAA) both increased voriconazole susceptibility would be consistent with this segment of the deletion overlapping a region important in azole resistance. Further experiments will be required to determine the precise role of the amino acids within the 855-879 region of AtrR.

The most striking finding of our analysis of the C-terminus of AtrR was our ability to separate the activation domains of this factor into two overlapping regions (Figure 7). The largest (from position 655 to 895 is required for normal transcriptional activation of *abcG1*. Any deletion in this region interfered with normal *abcG1* expression. Contained within this region was a smaller segment (655 to 755) that was required for normal *cyp51A* expression. We previously described the identification of a coactivator protein called NcaA that was required for normal *abcG1* expression but was dispensable for *cyp51A* transcription (21). This earlier finding is consistent with the larger region of AtrR required for *abcG1* expression relative to that for *cyp51A*. We suggest that the NcaA interaction region of AtrR may be between 755 and 895 and are carrying out experiments to test this suggestion.

**Figure 7.**
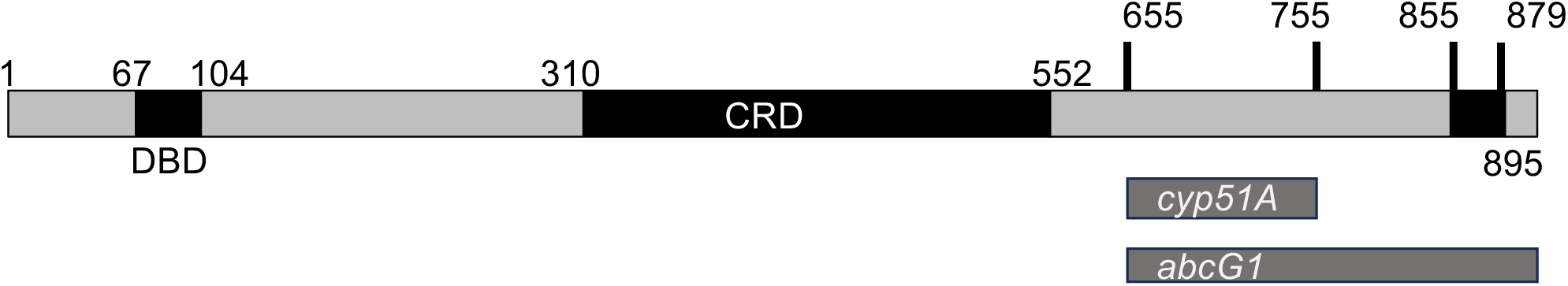
Functional domains in the AtrR transcription factor. The numbers refer to positions along the protein chain from *A. fumigatus*. The abbreviations for the identified protein regions are DNA-binding domain (DBD) and central regulatory domain (CRD). The minimal transcriptional activation domains required for normal expression of either *cyp51A* or *abcG1* are shown as gray bars.

Finally, the direct AtrR target genes that are reduced in expression by the presence of the Δ855-879 AtrR represent integral membrane proteins, with several thought to be transporters involved in nitrogen homeostasis. These include an ammonium transporter (Afu1g1930), two nitrate transporters (Afu1g12850, Afu1g17470) and a polyamine transporter (Afu5g14660). This is interesting in light of previous earlier ChIP-seq data that detected a nitrite reductase gene (niiA) as a target of AtrR and suggests a possible role for this transcription factor in the response to changes in nitrogen source (15).

Based on our RT-qPCR analysis, we expected to find *abcG1* as a target gene responsive to the presence of the Δ855-879 AtrR. The detection of the ABC transporter-encoding gene *mdr1* was less predictable but is consistent with data from several groups that have implicated this ABC transporter in azole resistance in *A. fumigatus* (22–26). We also detected AtrR binding to the *mdr1* promoter in an earlier ChIP-seq experiment (15). At least two additional transcription factors appear to act to regulate *mdr1*, Mdu2(OdrA) (24, 26) and SltA (25). While AtrR impacts expression of both *abcG1* and *mdr1*, Mdu2 and SltA effects are restricted to *mdr1*. Interestingly, another membrane transporter gene called *ena1* is dependent on AtrR and SltA. This gene encodes a sodium transporter and homologues of Ena1 have been associated with virulence defects and increased azole susceptibility in the pathogen *Cryptococcus neoformans* (27, 28). We are currently testing *ena1* mutants for their impact on azole drug resistance. Together, these data illustrate the complex transcriptional circuitry that regulates azole susceptibility in *A. fumigatus*. Our identification of the 855-879 region of AtrR as a contributor to transcription of membrane proteins that modulate azole susceptibility but surprisingly not a critical contributor to expression of *cyp51A* or other ergosterol biosynthetic enzyme-encoding genes. These data reveal the gene-specific regulation of azole susceptibility by the AtrR factor.

## Materials and Methods

### Strains and growth conditions

All strains used in this study were derived from the AfS35 strain and are listed in Supplemental Table 1. *A. fumigatus* strains were typically grown at 37°C in YG medium (0.5% Yeast extract and 2% Glucose). Selection of transformants was accomplished using minimal medium (MM; 1% glucose, nitrate salts, trace element mix, 2% agar [pH 6.5]). The trace element mix, vitamins, and nitrate salts are as described in the appendix of reference (Kafer E, 1977) supplemented with 1% sorbitol and 20 mg/liter phleomycin (InvivoGen) after adjusting the pH to 7.0. For solid medium, 1.5% agar was added.

### Plasmids

All plasmids used in this work are listed in Supplemental Table 2. The different *atrR* mutants were generated as follows: Plasmid pSR28 was constructed using Gibson assembly of 4 PCR fragments in the vector pUC19: 198 bp corresponding to the carboxy-terminus of the *atrR* gene, the transcription terminator of *cyc1*, the *ble* resistance marker gene, and a 45-bp region downstream of the *atrR* gene. The plasmid pSR28 was sequenced to confirm the desired construction and used as parent for construction of other plasmids (Chemically synthesized in this backbone by GenScript, USA) having internal and C-terminal deletions in the *atrR* gene (Table 2). For the alanine scanning mutagenesis, plasmids were generated that contained multiple substitution mutation that led to replace of 3 adjacent codons with alanine codons and introduced a SacII restriction site for screening purposes (Table 2). The *atrR* insert of these plasmids was released by KpnI and SalI restriction enzyme digestions and transformed into AfS35. Targeted integration of these strains was verified by PCR diagnosis of the novel junctions and sequence verified.

### Radial growth assay

Fresh spores of appropriate *A. fumigatus* strains were suspended in 1X phosphate-buffered saline (PBS) supplemented with 0.01% Tween 20 (1X PBST). The spore suspension was counted using a hemocytometer to determine the spore concentration. Spores were then appropriately diluted in 1X PBST. ∼100 spores (in 4 µl) were spotted on minimal medium with or without the drug. The plates were incubated at 37°C and inspected for growth every 12 h.

### Real-time PCR

Reverse transcription-quantitative PCR (RT-qPCR) was performed as described in (29) with the following modification. Each strain was inoculated in petri dishes containing 20 mL of YG broth at 37°C for 24 h (uninduced condition) and 16 h growth in YG, plus 8 h with voriconazole added YG medium (induced condition) under unshaking condition.

### Western blotting

Western blotting was performed as described in (29). The rabbit AtrR polyclonal antibody was used at a 1:1600 dilution.

### RNA-Sequencing

Approximately 10^6^ spores of each strain were inoculated in petri dishes containing 20 mL of YG broth at 37°C for 24 h (uninduced condition) and 16 h growth in YG then treated with voriconazole for 8 h (induced condition). Mycelia that formed as a biofilm on the top were collected (∼100 mg) and ground into a fine powder in liquid nitrogen using a mortar and pestle. Total RNA was then isolated from the ground mycelium using the Trizol reagent (Ambion) according to the manufacturer’s instructions. Traces of contaminating genomic DNA were eliminated by treating the RNA with 10X-turbo DN*ase* followed by a DN*ase* inactivation reagent (both reagents from Invitrogen).

RNA quality assessment was done by nanodrop and Tapestation (Genohub Inc., Texas, USA). Sequencing libraries preparation was done using NEBNext^®^ Ultra^TM^ II Directional RNA Library Prep Kit for Illumina (Genohub Inc., Texas, USA). Bioinformatics, utilizing standard methods, were carried out as previously described (30).

## Acknowledgments

This work was supported by NIH grants AI143198 and AI162802. We thank Drs. Damian Krysan, Scott Filler, Mike Bromley and Paul Bowyer for helpful discussions and Dr. Sanjoy Paul for a critical evaluation of this manuscript.

## Supplemental data

**Supplemental Figure 1.**
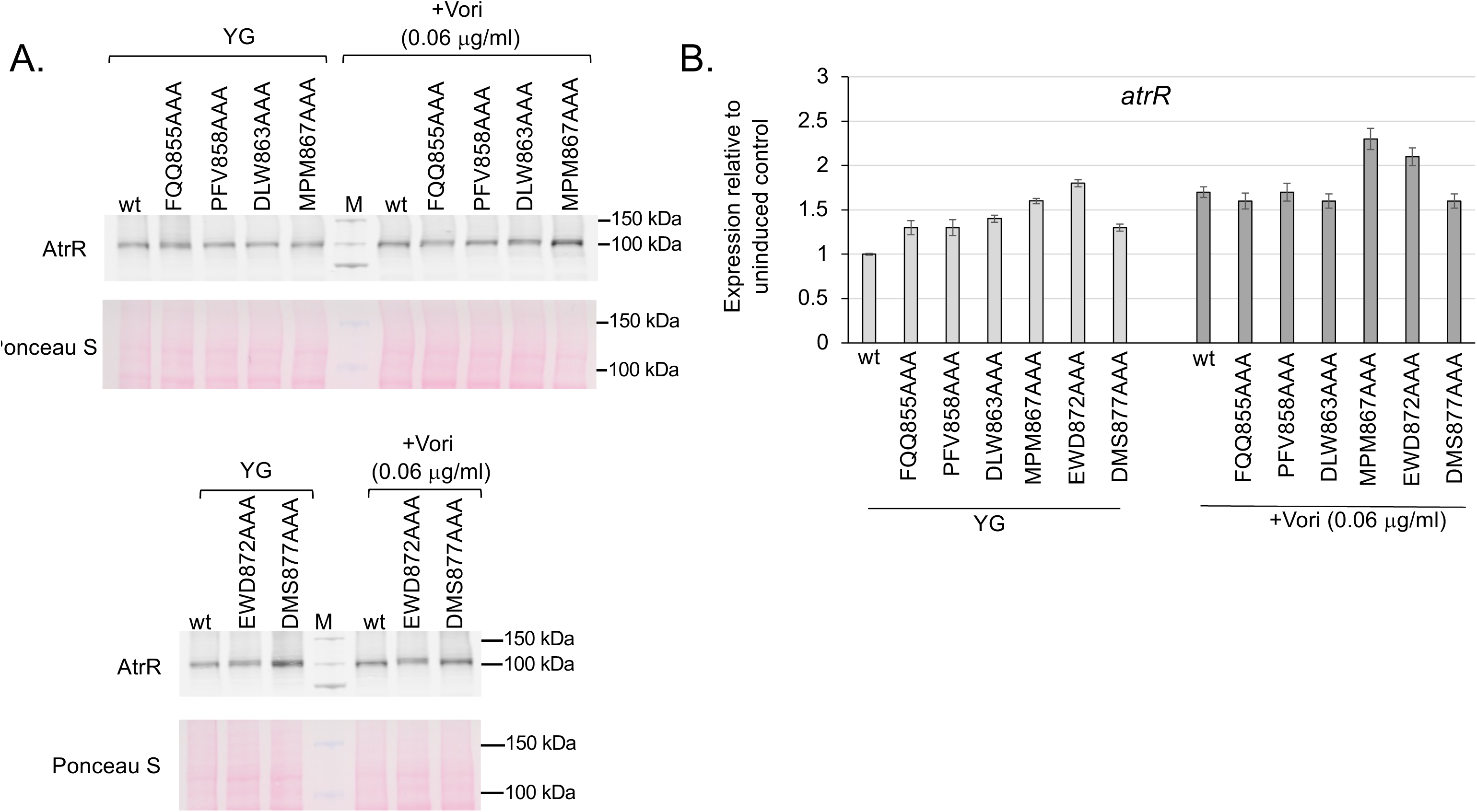
Equivalent expression of alanine-scanning mutants. A. Strains expressing the indicated alanine-scanning mutant forms of AtrR were analyzed by western blotting with the anti-AtrR antibody as described in the text. Molecular mass markers are indicated to the right side of each panel (M, Marker proteins). B. Levels of atrR mRNA for each strain were assessed using RT-qPCR. The level of each mRNA was normalized to that seen for the wild-type *atrR* gene grown in the absence of voriconazole.

**Supplemental table 1.** Strains used in this work.

| Strain Number. | Strain Name | Parent | Genotype | Reference |
| --- | --- | --- | --- | --- |
| 1 | AfS35 | D141 | *akuAΔ::loxP* | FGSC |
| 2 | SRF59 | AfS35 | *atrR::ble* | This study |
| 3 | SRF60 | AfS35 | FQQ855AAA<br>*atrR::ble* | This study |
| 4 | SRF61 | AfS35 | PFV858AAA<br>*atrR::ble* | This study |
| 5 | SRF62 | AfS35 | DLW863AAA<br>*atrR::ble* | This study |
| 6 | SRF63 | AfS35 | MPM867AAA<br>*atrR::ble* | This study |
| 7 | SRF64 | AfS35 | EWD872AAA<br>*atrR::ble* | This study |
| 8 | SRF65 | AfS35 | DMS877AAA<br>*atrR::ble* | This study |
| 9 | SRF66 | AfS35 | Δ855-879<br>*atrR::ble* | This study |
| 10 | SRF67 | AfS35 | 1-654 *atrR::ble* | This study |
| 11 | SRF68 | AfS35 | 1-754 *atrR::ble* | This study |
| 12 | SRF69 | AfS35 | 1-854 *atrR::ble* | This study |
| 13 | SRF70 | AfS35 | Δ655-754<br>*atrR::ble* | This study |
| 14 | SRF71 | AfS35 | Δ655-854<br>*atrR::ble* | This study |
| 15 | SPF323 | AfS35 | *atrRΔ::hph* | This study |

**Supplemental table 2.**
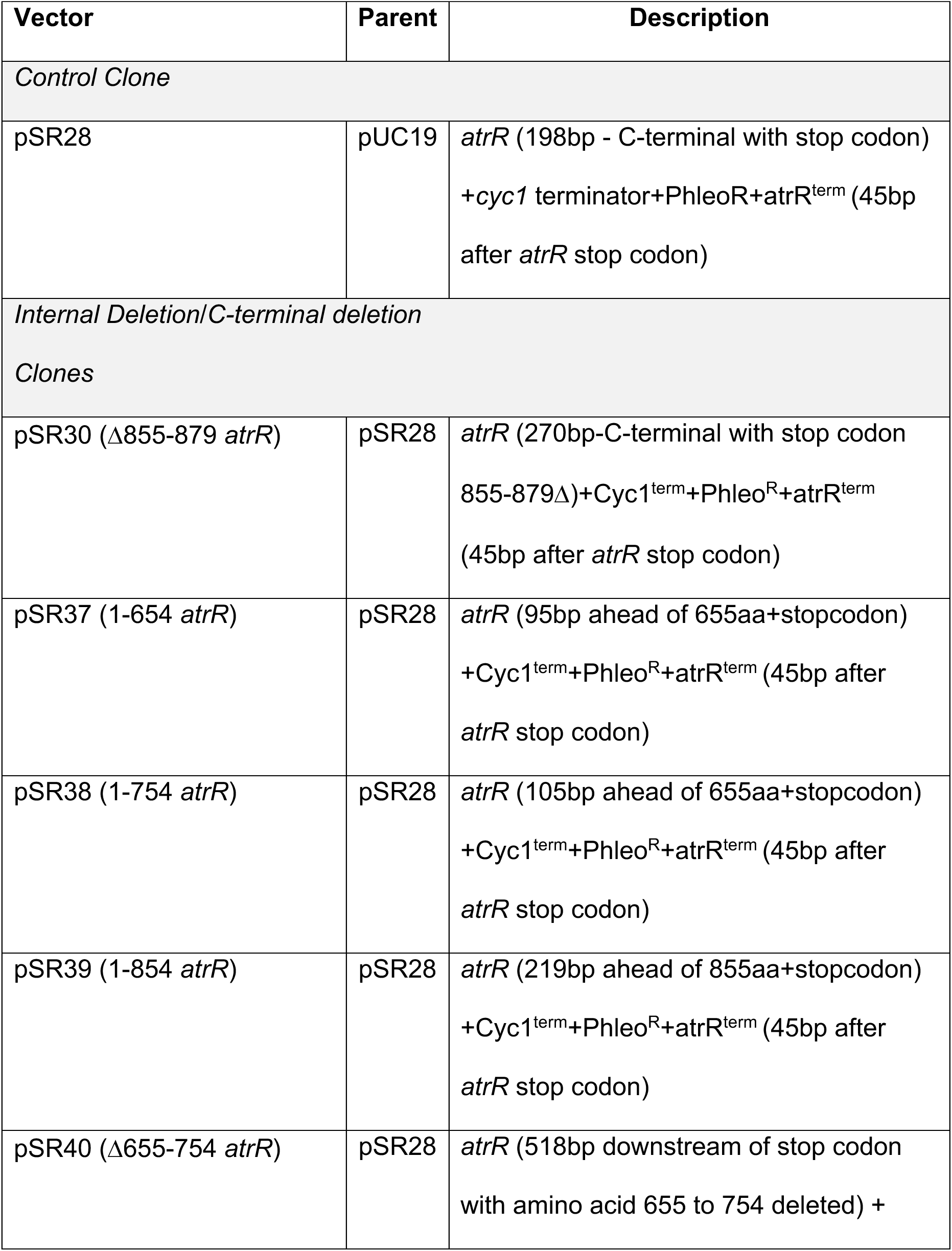
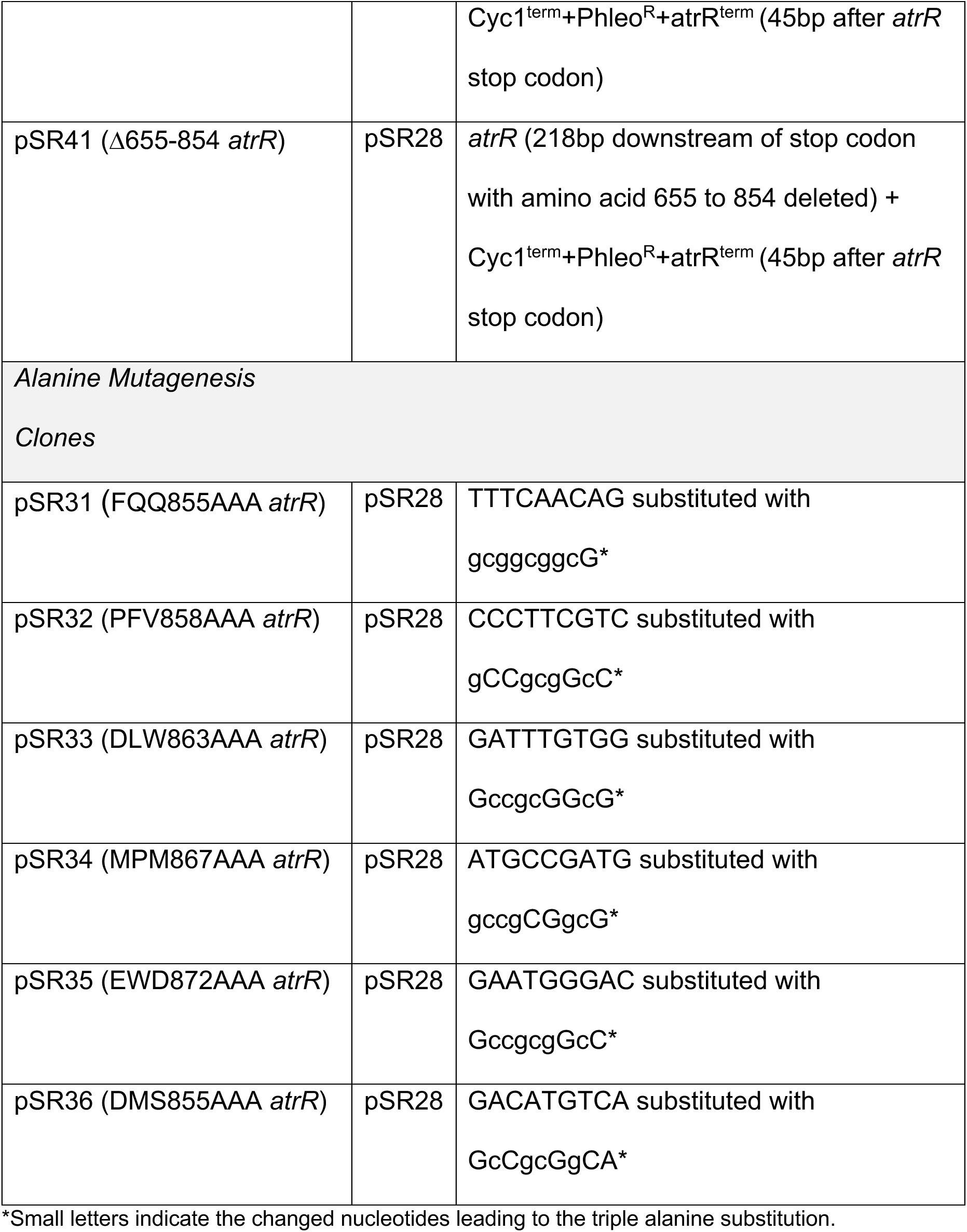
Plasmids used in this work.

## Literature cited

1. Lockhart SR, Chowdhary A, Gold JAW. 2023. The rapid emergence of antifungal-resistant human-pathogenic fungi. Nat Rev Microbiol doi:10.1038/s41579-023-00960-9.

2. Verweij PE, Snelders E, Kema GH, Mellado E, Melchers WJ. 2009. Azole resistance in Aspergillus fumigatus: a side-effect of environmental fungicide use? Lancet Infect Dis 9:789–95.

3. Snelders E, Melchers WJ, Verweij PE. 2011. Azole resistance in Aspergillus fumigatus: a new challenge in the management of invasive aspergillosis? Future Microbiol 6:335–47.

4. Verweij PE, Chowdhary A, Melchers WJ, Meis JF. 2016. Azole Resistance in Aspergillus fumigatus: Can We Retain the Clinical Use of Mold-Active Antifungal Azoles? Clin Infect Dis 62:362–8.

5. Rhodes J, Abdolrasouli A, Dunne K, Sewell TR, Zhang Y, Ballard E, Brackin AP, van Rhijn N, Chown H, Tsitsopoulou A, Posso RB, Chotirmall SH, McElvaney NG, Murphy PG, Talento AF, Renwick J, Dyer PS, Szekely A, Bowyer P, Bromley MJ, Johnson EM, Lewis White P, Warris A, Barton RC, Schelenz S, Rogers TR, Armstrong-James D, Fisher MC. 2022. Population genomics confirms acquisition of drug-resistant Aspergillus fumigatus infection by humans from the environment. Nat Microbiol 7:663–674.

6. Shelton JMG, Rhodes J, Uzzell CB, Hemmings S, Brackin AP, Sewell TR, Alghamdi A, Dyer PS, Fraser M, Borman AM, Johnson EM, Piel FB, Singer AC, Fisher MC. 2023. Citizen science reveals landscape-scale exposures to multiazole-resistant Aspergillus fumigatus bioaerosols. Sci Adv 9:eadh8839.

7. Snelders E, van der Lee HA, Kuijpers J, Rijs AJ, Varga J, Samson RA, Mellado E, Donders AR, Melchers WJ, Verweij PE. 2008. Emergence of azole resistance in Aspergillus fumigatus and spread of a single resistance mechanism. PLoS Med 5:e219.

8. Kuipers S, Bruggemann RJ, de Sevaux RG, Heesakkers JP, Melchers WJ, Mouton JW, Verweij PE. 2011. Failure of posaconazole therapy in a renal transplant patient with invasive aspergillosis due to Aspergillus fumigatus with attenuated susceptibility to posaconazole. Antimicrob Agents Chemother 55:3564–6.

9. van der Linden JW, Camps SM, Kampinga GA, Arends JP, Debets-Ossenkopp YJ, Haas PJ, Rijnders BJ, Kuijper EJ, van Tiel FH, Varga J, Karawajczyk A, Zoll J, Melchers WJ, Verweij PE. 2013. Aspergillosis due to voriconazole highly resistant Aspergillus fumigatus and recovery of genetically related resistant isolates from domiciles. Clin Infect Dis 57:513–20.

10. Willger SD, Puttikamonkul S, Kim KH, Burritt JB, Grahl N, Metzler LJ, Barbuch R, Bard M, Lawrence CB, Cramer RA, Jr. 2008. A sterol-regulatory element binding protein is required for cell polarity, hypoxia adaptation, azole drug resistance, and virulence in Aspergillus fumigatus. PLoS Pathog 4:e1000200.

11. Gsaller F, Hortschansky P, Furukawa T, Carr PD, Rash B, Capilla J, Muller C, Bracher F, Bowyer P, Haas H, Brakhage AA, Bromley MJ. 2016. Sterol Biosynthesis and Azole Tolerance Is Governed by the Opposing Actions of SrbA and the CCAAT Binding Complex. PLoS Pathog 12:e1005775.

12. Paul S, Verweij PE, Melchers WJG, Moye-Rowley WS. 2022. Differential Functions of Individual Transcription Factor Binding Sites in the Tandem Repeats Found in Clinically Relevant cyp51A Promoters in Aspergillus fumigatus. mBio 13:e0070222.

13. Kuhbacher A, Peiffer M, Hortschansky P, Merschak P, Bromley MJ, Haas H, Brakhage AA, Gsaller F. 2022. Azole Resistance-Associated Regulatory Motifs within the Promoter of cyp51A in Aspergillus fumigatus. Microbiol Spectr 10:e0120922.

14. Hagiwara D, Miura D, Shimizu K, Paul S, Ohba A, Gonoi T, Watanabe A, Kamei K, Shintani T, Moye-Rowley WS, Kawamoto S, Gomi K. 2017. A Novel Zn2-Cys6 Transcription Factor AtrR Plays a Key Role in an Azole Resistance Mechanism of Aspergillus fumigatus by Co-regulating cyp51A and cdr1B Expressions. PLoS Pathog 13:e1006096.

15. Paul S, Stamnes M, Thomas GH, Liu H, Hagiwara D, Gomi K, Filler SG, Moye-Rowley WS. 2019. AtrR Is an Essential Determinant of Azole Resistance in Aspergillus fumigatus. MBio 10.

16. MacPherson S, Larochelle M, Turcotte B. 2006. A fungal family of transcriptional regulators: the zinc cluster proteins. Microbiol Mol Biol Rev 70:583–604.

17. Sang H, Hulvey JP, Green R, Xu H, Im J, Chang T, Jung G. 2018. A Xenobiotic Detoxification Pathway through Transcriptional Regulation in Filamentous Fungi. mBio 9.

18. Schoen TJ, Calise DG, Bok JW, Giese MA, Nwagwu CD, Zarnowski R, Andes D, Huttenlocher A, Keller NP. 2023. Aspergillus fumigatus transcription factor ZfpA regulates hyphal development and alters susceptibility to antifungals and neutrophil killing during infection. PLoS Pathog 19:e1011152.

19. Hahn S, Young ET. 2011. Transcriptional regulation in Saccharomyces cerevisiae: transcription factor regulation and function, mechanisms of initiation, and roles of activators and coactivators. Genetics 189:705–36.

20. Ferrari S, Ischer F, Calabrese D, Posteraro B, Sanguinetti M, Fadda G, Rohde B, Bauser C, Bader O, Sanglard D. 2009. Gain of function mutations in CgPDR1 of Candida glabrata not only mediate antifungal resistance but also enhance virulence. PLoS Pathog 5:e1000268.

21. Paul S, Ror S, McDonald WH, Moye-Rowley WS. 2022. Biochemical Identification of a Nuclear Coactivator Protein Required for AtrR-Dependent Gene Regulation in Aspergillus fumigatus. mSphere 7:e0047622.

22. Nascimento AM, Goldman GH, Park S, Marras SA, Delmas G, Oza U, Lolans K, Dudley MN, Mann PA, Perlin DS. 2003. Multiple resistance mechanisms among Aspergillus fumigatus mutants with high-level resistance to itraconazole. Antimicrob Agents Chemother 47:1719–26.

23. Fraczek MG, Bromley M, Buied A, Moore CB, Rajendran R, Rautemaa R, Ramage G, Denning DW, Bowyer P. 2013. The cdr1B efflux transporter is associated with non-cyp51a-mediated itraconazole resistance in Aspergillus fumigatus. J Antimicrob Chemother doi:10.1093/jac/dkt075.

24. Sturm L, Geissel B, Martin R, Wagener J. 2020. Differentially Regulated Transcription Factors and ABC Transporters in a Mitochondrial Dynamics Mutant Can Alter Azole Susceptibility of Aspergillus fumigatus. Front Microbiol 11:1017.

25. Du W, Zhai P, Wang T, Bromley MJ, Zhang Y, Lu L. 2021. The C(2)H(2) Transcription Factor SltA Contributes to Azole Resistance by Coregulating the Expression of the Drug Target Erg11A and the Drug Efflux Pump Mdr1 in Aspergillus fumigatus. Antimicrob Agents Chemother 65.

26. Sasse C, Bastakis E, Bakti F, Hofer AM, Zangl I, Schuller C, Kohler AM, Gerke J, Krappmann S, Finkernagel F, Harting R, Strauss J, Heimel K, Braus GH. 2023. Induction of Aspergillus fumigatus zinc cluster transcription factor OdrA/Mdu2 provides combined cellular responses for oxidative stress protection and multiple antifungal drug resistance. mBio 14:e0262823.

27. Idnurm A, Walton FJ, Floyd A, Reedy JL, Heitman J. 2009. Identification of ENA1 as a virulence gene of the human pathogenic fungus Cryptococcus neoformans through signature-tagged insertional mutagenesis. Eukaryot Cell 8:315–26.

28. Jung KW, Strain AK, Nielsen K, Jung KH, Bahn YS. 2012. Two cation transporters Ena1 and Nha1 cooperatively modulate ion homeostasis, antifungal drug resistance, and virulence of Cryptococcus neoformans via the HOG pathway. Fungal Genet Biol 49:332–45.

29. Paul S, Diekema D, Moye-Rowley WS. 2017. Contributions of both ATP-Binding Cassette Transporter and Cyp51A Proteins Are Essential for Azole Resistance in Aspergillus fumigatus. Antimicrob Agents Chemother 61.

30. Paul S, Stamnes MA, Moye-Rowley WS. 2023. Interactions between the transcription factors FfmA and AtrR are required to properly regulate gene expression in the fungus Aspergillus fumigatus. G3 (Bethesda) 13.

